# K2R: Tinted de Bruijn Graphs implementation for efficient read extraction from sequencing datasets

**DOI:** 10.1101/2024.02.15.580442

**Authors:** Léa Vandamme, Bastien Cazaux, Antoine Limasset

## Abstract

The analysis of biological sequences often depends on reference genomes; however, achieving accurate assemblies remains a significant challenge. As a result, de novo analysis directly from raw sequencing reads, without pre-processing, is frequently a more practical approach. A common need across various applications is the ability to identify reads containing a specific *k*-mer within a dataset. This *k*-mer-to-read association is critical in multiple contexts, such as genotyping, bacterial strain resolution, profiling, data compression, error correction, and assembly. While this challenge appears similar to the extensively researched colored de Bruijn graph problem, resolving it at the read level is prohibitively resource-intensive for practical applications. In this work, we demonstrate its tractable resolution by leveraging reasonnable assumptions for genome sequencing dataset indexing. To tackle this challenge, we introduce the Tinted de Bruijn Graph concept, an altered version of the colored de Bruijn graph where each read in a sequencing dataset acts as a distinct source. We developed K2R, a highly scalable index that implements this framework efficiently. K2R’s performance, in terms of index size, memory footprint, throughput, and construction time, is benchmarked against leading methods, including hashing techniques (e.g., Short Read Connector and Fulgor), full-text indexing (e.g., Movi and Themisto) across various datasets. To demonstrate K2R’s scalability, we indexed two human datasets from the T2T consortium. The 126X coverage ONT dataset was indexed in 9 hours using 61GB of RAM, resulting in a 30GB index. Similarly, the 56X coverage HiFi dataset was indexed in less than 5 hours using 39GB of RAM, producing a 20.5GB index. Developed in C++, the K2R index is open-source and available on GitHub at http://github.com/LeaVandamme/K2R.

## Introduction

The pursuit of direct and unbiased access to complete nucleic sequences remains a central focus in sequence analysis. Despite significant advancements over the past two decades, achieving accurate and comprehensive reference sequences continues to pose major challenges (He et al. 2023). A primary difficulty lies in the inherent complexity of assembly processes, both from a theoretical perspective (Nagarajan and Pop 2009) and in practical implementation (Nurk et al. 2022). These challenges are often exacerbated by the intrinsic properties of sequencing data, which can limit the direct applicability of assemblies to key biological questions, such as quantification or dataset comparison. Consequently, the *de novo* analysis of raw datasets holds paramount importance.

Since the introduction of the seed-and-extend paradigm (Altschul et al. 1990), numerous applications have heavily relied on identifying fixed-length word matches, commonly referred to as *k-mers*, to detect sequence similarities. A highly efficient representation of a *k*-mer set from sequencing datasets is the de Bruijn graph (Chikhi et al. 2015), which partially preserves the original ordering of *k*-mers. *K*-mer sets have found widespread applications across diverse domains, including quantification (Wang et al. 2012), dataset comparison (Ramos et al. 2022), phylogeny (Lyman et al. 2017), assembly (Bankevich et al. 2012, 2022), compression (Benoit et al. 2015), error correction (Limasset et al. 2020), and sequence alignment (Liu et al. 2016; Heydari et al. 2018). The advent of short-read sequencing technologies has further highlighted the utility of the de Bruijn graph as a versatile and efficient structure for handling *k*-mers. Consequently, *k*-mer indexes have become well-researched data structures, with highly efficient implementations leveraging either hashing-based approaches (Marchet et al. 2021b; Pibiri 2022) or full-text indexing methods (Alanko et al. 2023; Rossi et al. 2022).

A significant limitation of such indexes is their inability to retain critical information, specifically the colocalization of *k*-mers within shared DNA fragments. In the context of short reads, this loss of information was considered negligible, as the *k*-mer size (*k*) typically approximated the size of the reads. However, for long reads, this limitation becomes far more impactful, as it results in the omission of highly valuable long-range information (Hu et al. 2021). One potential solution is the ability to identify the specific reads in which a given *k*-mer appears. Existing tools based on document scanning, while practical, have the drawback of requiring the entire, often redundant, dataset to be read for extracting the relevant reads, leading to very low throughput (Baire et al. 2024). In contrast, full-text indexes (Li 2014), which can efficiently locate patterns within indexed datasets, present a natural and more effective alternative. Recent advancements in BWT-based full-text indexes have led to the development of two novel structures that refine the previously dominant FM-index. The r-index (Bannai et al. 2020), a full-text index requiring space proportional to *𝒪*(*r*), where *r* denotes the number of BWT runs for an input text of size *n*, introduces a suffix array sampling strategy that occupies only 2*r* space. This innovation significantly reduces locate time from Ω(*n/r*) per occurrence (as in traditional FM-indexes) to *𝒪*(log(*n/r*)). More recently, an extension of the r-index, br-index (Arakawa et al. 2022) (bi-directional index) has been developped enabling bidirectional extensions along the pattern search process. This use *𝒪*(*r* + *r*_*R*_) words of space, where *r*_*R*_ is the number of BWT runs of the reversed text, and the locate time is *𝒪*(*occ*), where *occ* is the number of occurences of a pattern in the text. Movi (Zakeri et al. 2023), based on the move index (Nishimoto and Tabei 2020), achieves both *𝒪*(*r*) space efficiency and *𝒪*(1)-time queries. Nonetheless, these structures face challenges in compressing the high redundancy of reads, which often contain noise that disrupts redundancy and introduces irrelevant novel sequences to index.

This *k*-mer-to-reads problem closely resembles the well-studied colored de Bruijn graph challenge (Iqbal et al. 2012). A colored de Bruijn graph, constructed from a collection of documents, associates each *k*-mer with the list of documents in which it appears. In principle, each read could be treated as an individual document to leverage existing colored de Bruijn graph implementations. However, this approach becomes prohibitively expensive given the vast number of reads anticipated from gigabase level genomes sequencing, easily reaching tens of millions when indexing ten thousands dataset is already very expensive (Marchet and Limasset 2023; Marchet et al. 2021a). Despite this, we emphasize that assumptions about the *k*-mer distribution in colored de Bruijn graphs are often limited. In contrast, we aim to demonstrate the problem tractability of indexing a sequencing dataset by leverage realistic properties.

In this work, we examine these properties and introduce practical techniques to leverage them effectively. For practical purposes, we introduce a scalable and efficient tool named K2R (*K*-mer to Reads), which leverages sequencing datasets properties, and perform an extensive benchmark against state-of-the-art structures capable of executing similar tasks. The K2R index, developed in C++, is open source and available on Github http://github.com/LeaVandamme/K2R.

## Results

All experiments were conducted on a single cluster node equipped with an Intel(R) Xeon(R) Gold 6130 CPU @ 2.10GHz, 128GB of RAM, and running Ubuntu 22.04. To validate our results on real data, we chose to perform the same analysis on closely related *E*.*Coli* sequencings. Specifically, we selected a recent ONT sequencing with a low error rate, approximately 3% (Accession SRR26899125), a 20kb HiFi sequencing (Accession SRR11434954), and a HiSeq X Ten paired-end sequencing (Accession DRR395239). To extend our evaluation to a medium-sized genome, we included two *C*.*Elegans* datasets, one ONT sequencing (Accession SRR24201716) and another HiSeq X Ten paired-end sequencing (Accession ERR10914908).

Finally, to assess K2R scalability, we selected two human datasets from the T2T project. The HiFi datasets (SRX7897685, SRX7897686, SRX7897687, SRX7897688, and SRX5633451) together amount to 56.8X coverage, comprising both 20kb and 10kb libraries. The ONT sequencings, detailed at http://github.com/marbl/CHM13/blob/ master/Sequencing_data.md, total 126X coverage with an approximate error rate of 6%.

### State of the art

We categorize tools capable of processing Tinted de Bruijn graph queries into two main types. The first type includes full-text indexes, which can locate *k*-mer occurrences within an indexed read file. Utilizing a starting position array enables the determination of specific reads in which a queried *k*-mer is present. The second approach involves *k*-mer indexes, which rely on associative structures such as hash functions. Despite significant advancements in reducing the memory footprint of *k*-mer indexing to below 10 bits per *k*-mer (Pibiri 2022), a major challenge persists in managing the associated read identifier lists, as these lists are expected to be larger by several orders of magnitude.

Recent developments in BWT-based full-text indexes have introduced several innovative structures enhancing the capabilities of the FM index, which dominated the field for over a decade. The latest tool utilizing the r-index for large-scale queries is SPUmoni (Ahmed et al. 2023), while br-index (Arakawa et al. 2022) is an extension of the r-index and full-br-index a fully functional br-index. Movi (Zakeri et al. 2023) is the sole tool based on the move index. Given the efficiency of the move index, particularly in query time (*𝒪*(1)), we opted to include Movi in our benchmark over SPUmoni.

To our knowledge, the only specialized tools implementing *k*-mer to reads index is SRC (Marchet et al. 2020) employing Minimal Perfect Hash Function (Limasset et al. 2017; Pibiri and Trani 2021a) but we also included Themisto (Alanko et al. 2023) based on a Spectral BWT, Movi (Zakeri et al. 2023) based on the move index and Fulgor (Fan et al. 2023) based on Minimal Perfect Hash Functions (Pibiri and Trani 2021b) in our benchmark.

As outlined in the introduction, a fundamental limitation of both full-text and *k*-mer-based indexes is their difficulty in handling sequencing errors, leading to the indexing of irrelevant sequences. In contrast, *k*-mer abundance filtering enables *k*-mer indexing methods to effectively exclude non-essential *k*-mers by considering their frequency of occurrence.

In our benchmarking, both SRC and K2R indexed 31-mers while excluding those that appeared only once or more than 5,000 times in the dataset. This creates a theoretical disparity in comparisons, as full-text indexes inherently index the entire text, and Fulgor and Themisto do not offer a similar option. However, in practical terms, users of these techniques have no other options but to employ them as they currently exist, and most analyses actually filter out at least unique *k*-mers as they are considered unreliable. All minimizers-based methods in our benchmark use minimizers of length 15.

### Index Construction

We first evaluate the cost of constructing an index from whole genome sequencing datasets with the different tools.

In Figure 1, we assess the time and memory required according to the input coverage on three *E*.*Coli* datasets. We observe that for all tools, both time and memory requirements grow linearly with the input size. However, their respective performance varies greatly. Firstly, we omitted full-br-index from the benchmark since the smallest dataset provided (10X coverage of simulated reads from the *E*.*Coli* genome) took more than 24 minutes to process (about 20 times longer than Movi, 150 times longer than K2R). This suggests that full-br-index does not scale well for this application.

**Fig. 1.**
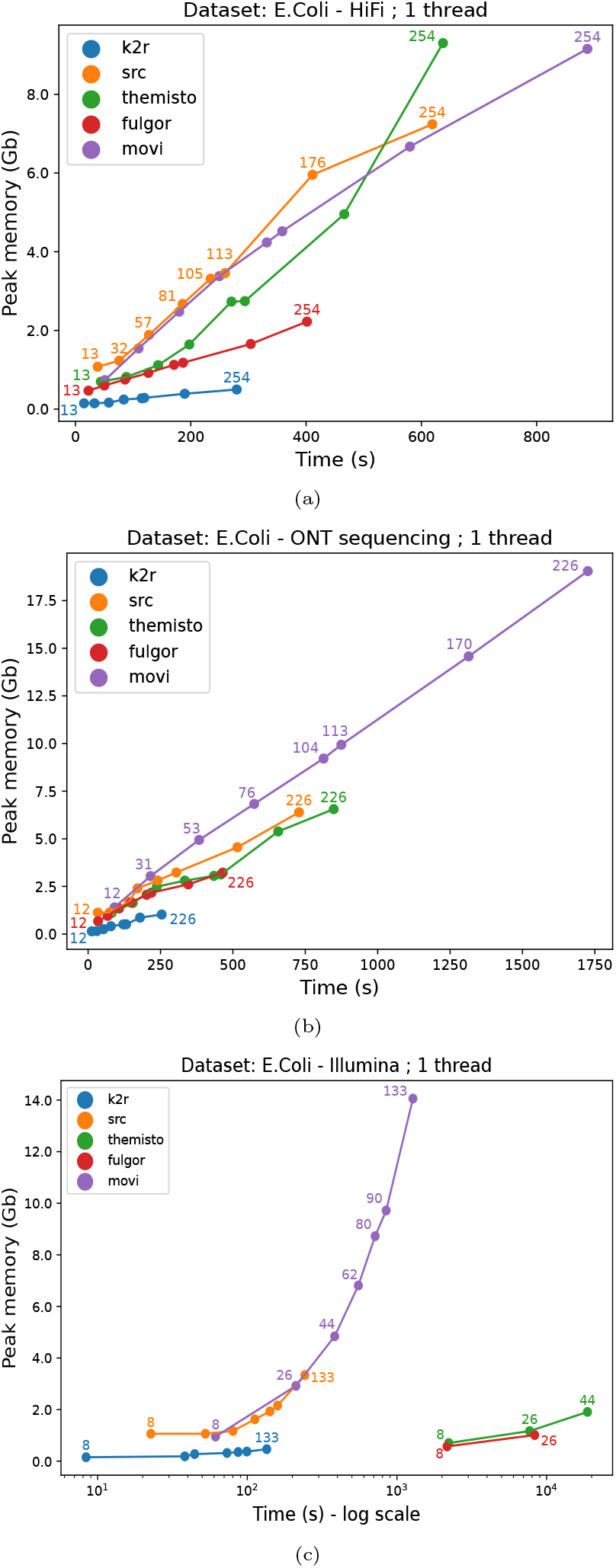
Memory peak and wall-clock time used during the index construction with three different kind of data with varying coverages (specified on the labels) from *E*.*Coli* genome : (a) HiFi (top), (b) ONT (center) and (c) Illumina (bottom) where we notice that both Themisto and Fulgor can’t scale to a 62X and 44X coverage respectively. This last graph is in logarithmic scale for readability purposes.

Overall, we observe that K2R is significantly faster and memory efficient that the state of the art, while Movi is slower and more memory expensive. This effect is quite small for HiFi data but is amplified with ONT data. For Illumina data, we note that the time required by Fulgor and Themisto increases dramatically. This is due to their operational mode, which requires a unique sequence per file. In our case, this means one file per read, leading to numerous disk accesses and a corresponding performance bottleneck.

We performed the same experiment on the *C*.*Elegans* genome, which is 20 times larger, as shown in Figure 2 and obtained similar results, highlighting K2R’s scalability in memory and, more importantly, in time. Since most of the memory is allocated to filter minimizers in a fixed size counting bloom filter (2^32^) across experiments, this explains K2R’s almost constant memory, even for larger datasets. As a comparison, we have added a curve using a smaller filter size (2^26^) that implies more minimizer collisions and potentially a higher number of false positives in the structure. We notice that Themisto and Fulgor does not appear for Illumina dataset. This is because these tools require a unique sequence per file, meaning that in our case, one file is created per read. These tools are better suited for indexing entire genomes, not datasets divided into individual reads.

**Fig. 2.**
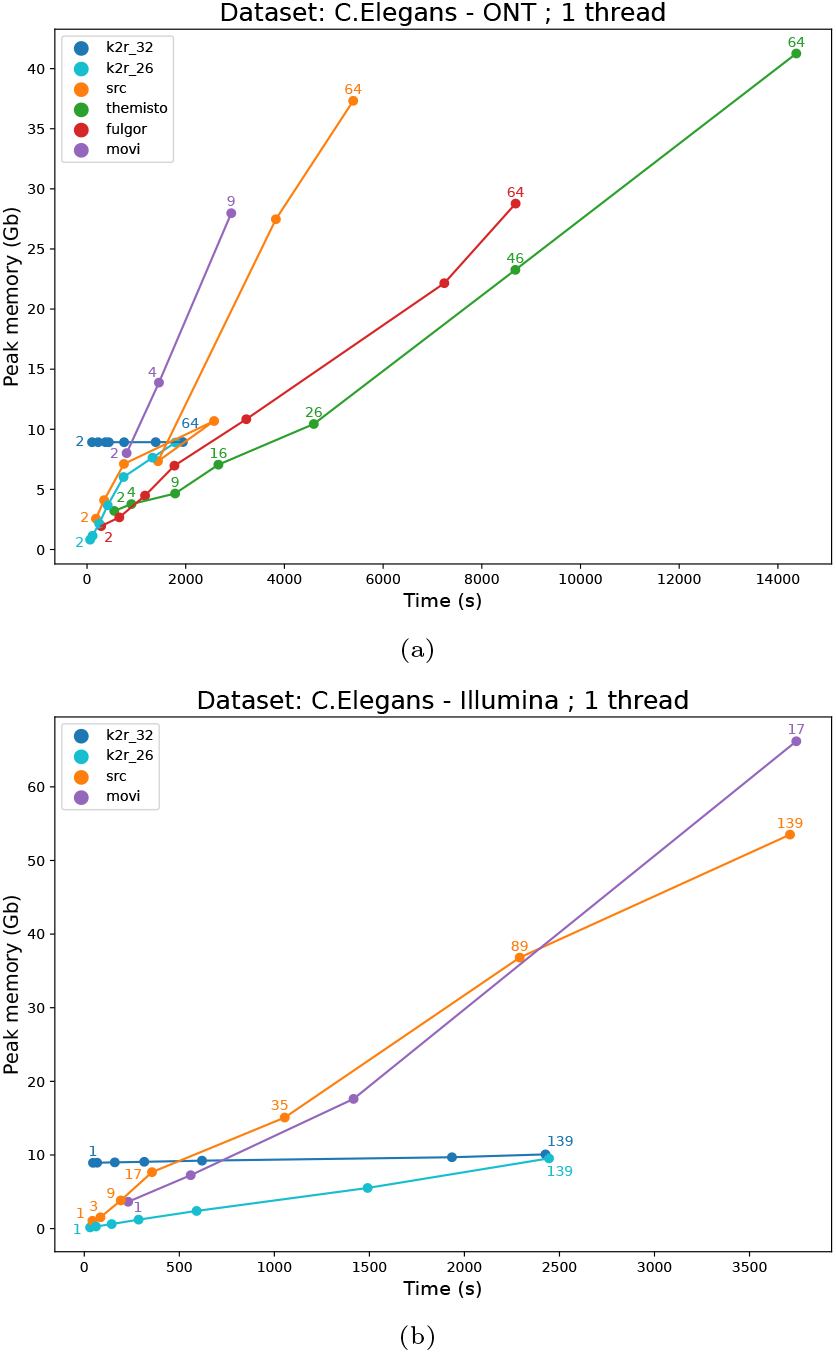
Memory peak and wall-clock time used during the index construction with two different kind of dataset from *C*.*Elegans* genome with varying coverages (specified on the labels) : (a) ONT (top), we notice that Movi can’t scale up for a coverage greater than 9X. (b) Illumina (bottom), we notice that Movi can’t scale up for a coverage greater than 17X. Themisto and Fulgor can’t be used with this type of data because of their mode of use. K2R 26 and K2R 32 refer to the counting bloom filter size (2^26^ or 2^32^) used to index.

**Fig. 3.**
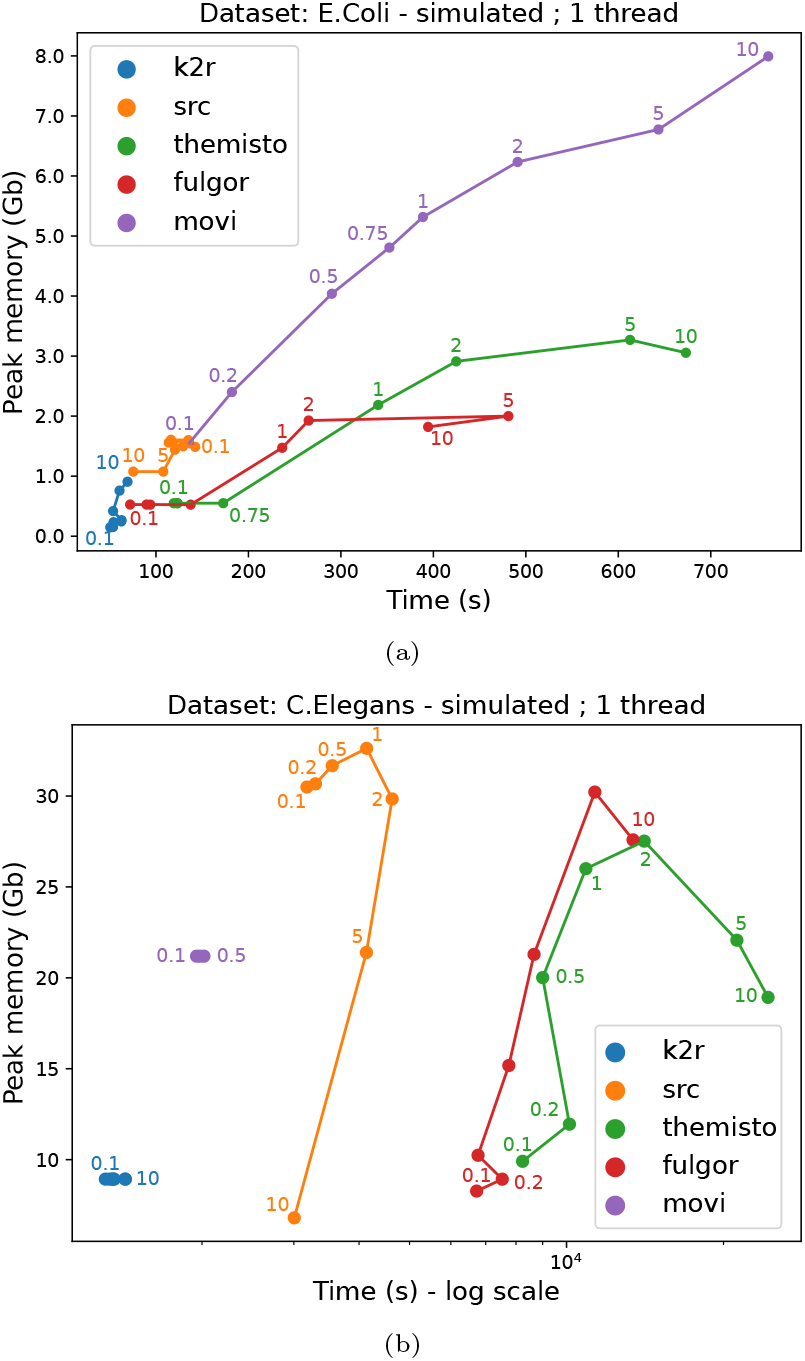
Memory peak and wall-clock time used during the index construction according to the error rate (specified on the labels in percent) for simulated reads (length of 10,000 with a 50X coverage) from (a) *E*.*Coli* (top) and (b) *C*.*Elegans* (bottom) genomes. This last graph is in logarithmic scale for readability purposes.

As a validation, we also performed the benchmark on simulated reads from the reference genomes, obtaining very similar results displayed in Figure S2 and S3 of the Appendix. The analysis was also conducted using multiple threads, as shown in Figures S5 and S6 of the Appendix.

Additionally, we applied this analysis to the two human datasets, using K2R exclusively due to scalability issues with other tools. We tested two values for maximum abundance (256 and 1,000). We note that this restriction is common and used by assembly tools like MECAT (Xiao et al. 2017) or aligners like BELLA (Guidi et al. 2018), skipping *k*-mers seen more than 128 times. The results, presented in Table 1, reveal the high cost of indexing highly repeated *k*-mers in complex genomes like that of humans. Including such *k*-mers nearly doubles both time and memory requirements.

**Table 1.** Index construction results on two *Human* datasets (56X HiFi and 126X ONT) using K2R, depending on the maximum abundance of minimizers. For each value, the RAM used, the wall-clock time and the structure size are presented in gigabytes and hours.

| Abundance | Max abundance: 256 |  | Max abundance: 1000 |  |
| --- | --- | --- | --- | --- |
| Dataset | HiFi 56X | ONT 126X | HiFi 56X | ONT 126X |
| RAM | 16 GB | 30 GB | 39 GB | 61 GB |
| Wall-clock time | 2h37 | 5h53 | 4h30 | 9h05 |
| Structure size | 7.6GB | 10.71GB | 20.5GB | 30.1GB |

These results demonstrate that such indexes are quite tractable to construct on various datasets, even on gigabase-scale genomes, on a modest workstation.

To further analyze the impact of noise, we evaluated in Figure 3 the time and memory requirements according to the input error rate, with coverage fixed at 50X. Since testing such parameters on real data poses challenges, this benchmark was conducted exclusively on simulated reads from the *E*.*Coli* and *C*.*Elegans* reference genomes. For the *E*.*Coli* data, this analysis revealed two distinct patterns. The error rate had minimal impact on the construction time for K2R and SRC, while it affected Fulgor, Movi, and Themisto. In the case of *C*.*Elegans*, the error rate has a minor influence on the memory usage of K2R and Movi (though Movi could not scale beyond an error rate of 0.5%). It can even decrease with a higher error rate as fewer *k*-mers are indexed due to abundance filtering. Full-text indexes, on the other hand, were significantly hindered by the error rate. They performed well with a low error rate but became much more costly with higher error rates. If SRC outperforms other methods in terms of time, regardless of the error rate, K2R itself is vastly superior in both terms of time and memory efficiency.

### Index Size

While the construction cost is a key factor in tractability, the actual final index size is more important since the construction needs to be performed only once but the index can be used multiple times. As before, we first assess the impact of coverage in Figure 4 for the *E*.*Coli* dataset and in Figure 5 for the *C*.*Elegans* datasets.

**Fig. 4.**
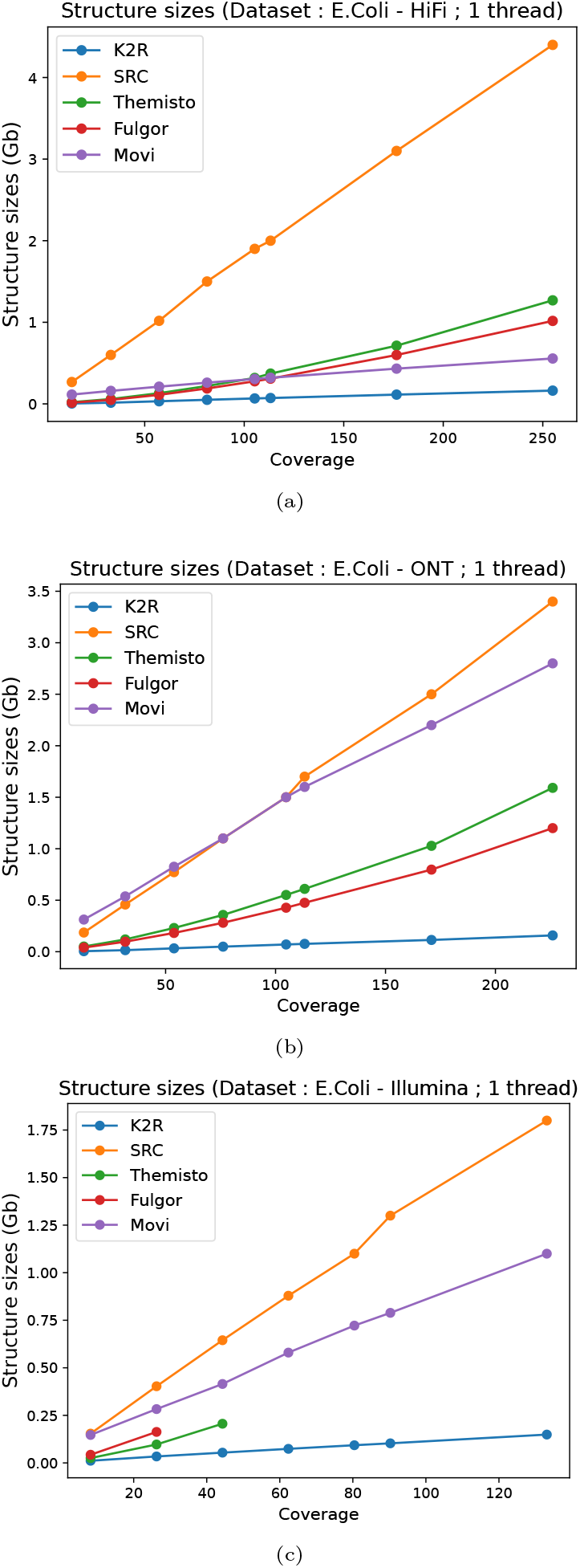
Comparison of the index size, for three different *E*.*Coli* genome datasets according to the input coverage: (a) HiFi dataset (top), (b) ONT dataset (center) and (c) Illumina dataset (bottom), where we notice that Themisto and Fulgor can’t scale up to coverages greater than 44 and 26 respectively.

**Fig. 5.**
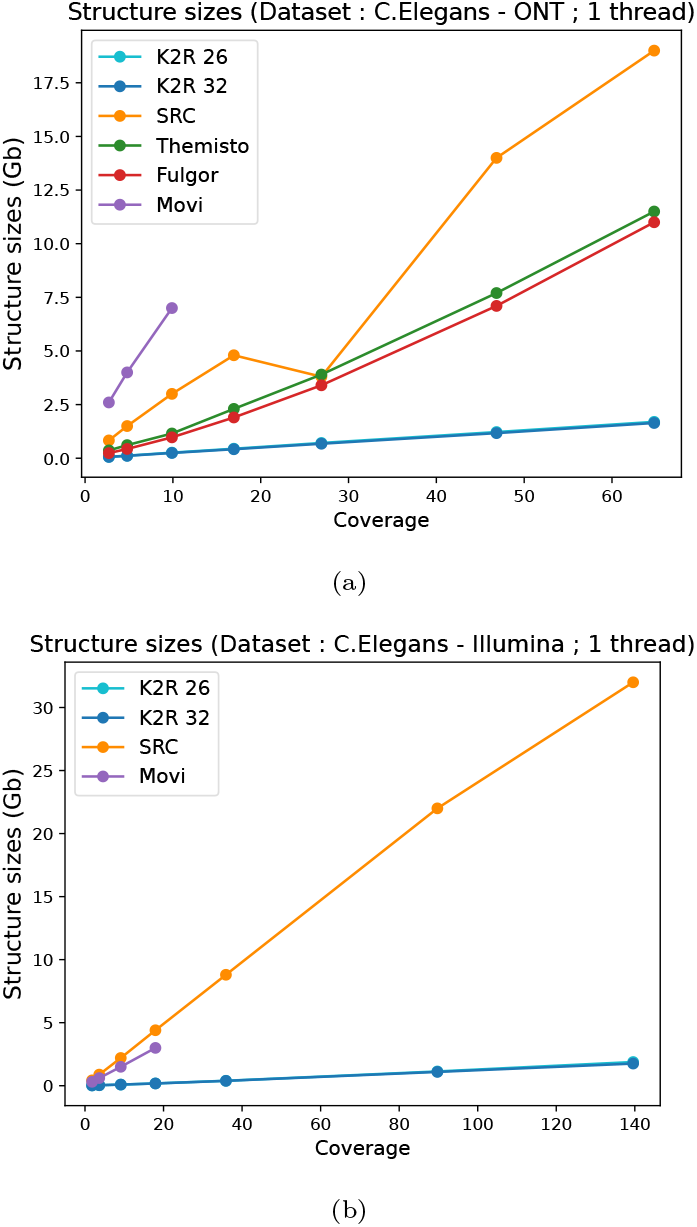
Comparison of the index size, for two types of dataset from *C*.*Elegans* genome according to the input coverages : (a) ONT and (b) Illumina. K2R 26 and K2R 32 refer to the counting bloom filter size (2^26^ or 2^32^) used to index, here the two curves are merged because the difference is almost non-existent.

We observe distinct patterns across the different data types. For HiFi data, all tools, except SRC, perform quite well using less or about a gigabyte for 200X coverage. However, for ONT data, only K2R maintains this efficiency, and Movi grows similarly to SRC. Illumina data follow the same trend but due to their usage mode, Themisto and Fulgor can’t be tested at high coverage levels. The differences between SRC/Movi and K2R are even more pronounced for the *C*.*Elegans* dataset. Notably, varying the counting Bloom filter size in K2R between 2^32^ and 2^26^ has minimal impact on the final structure size, as collisions remain negligible. To validate these results, we benchmarked simulated reads from the *E*.*Coli* and *C*.*Elegans* genomes, obtaining comparable outcomes (see Figures S2 and S3 in the Appendix). Finally, as a scalability experiment, we report the index size from human datasets (skipping *k*-mers seen more than 1,000 times). While indexing high-coverage ONT human datasets is relatively inexpensive in practice (about 30GB, which is more than an order of magnitude smaller than the actual dataset : 381GB), we find that the HiFi data index is even an order of magnitude smaller (20.5GB instead of 165GB), indicating that sequencing errors can significantly impact the size of such indexes (see Table 1).

Following the experimental plan, in a second experiment, we evaluate the impact of sequencing error rates on index size using simulated data, as presented in Figure S4 in the Appendix. As expected, a higher error rate generally means a larger index file. An exception is observed with SRC, where increased error rates reduce the number of colors indexed, leading to a smaller overall index size. Apart from this effect, K2R is almost always an order of magnitude smaller than its competitors, especially for larger datasets.

### Queries

We now assess the query throughput of the different indexes. Instead of measuring the query time for a single query (which is challenging due to overheads such as index loading), we tested batches of sequences ranging from 1,000 to 10,000, each between 500 and 10,000 bases long. We conducted two types of queries: positive queries, where reads were extracted from the *E. coli* reference genome, and negative queries, where reads were randomly generated. All queries were executed using 24 threads. The results are detailed in Figures S7 and S8 of the Appendix. It is important to note that Movi was excluded from these results, as it performs a different type of operation, focusing on presence/absence and similarity rather than returning the location of reads.

First, concerning the memory peak, K2R, Themisto and Fulgor are constant, while SRC grows quickly with the number of queries. This trend is consistent for both positive and negative queries.

In terms of time, however, we have different observations. SRC is still the less efficient (at least an order of magnitude slower than the others, two for 10,000 queries). Themisto follows, performing similarly to K2R for positive queries (around 10 seconds per experiment) but slightly slower for negative queries. K2R ranks next, with Fulgor emerging as the fastest, particularly for negative queries, and only marginally slower for a large number of positive queries. In practice, the time difference between K2R, Themisto, and Fulgor is minimal (less than 10 seconds in all cases), rendering all three tools, including K2R, highly efficient.

In summary, our analysis leads us to three key conclusions. Firstly, the K2R index stands out as the most efficient for construction, significantly reducing memory and time consumption, in stark contrast to the resource-intensive nature of BWT-based indexes. Secondly, and perhaps most critically, K2R indexes are markedly smaller than current leading tools, surpassing them by several orders of magnitude, independently of coverage and error rates. Thirdly, queries processed through K2R are fast and as memory frugal than Themisto and Fulgor.

These findings display the practical feasibility of employing Tinted de Bruijn graph methodologies at scale, highlighting the effectiveness of the presented approaches.

## Discussion

The K2R index holds significant potential for diverse applications requiring scalable read-level resolution, such as metagenomics or transcriptomics quantification, assembly, or genotyping. Future work will aim to integrate and fine-tune it for these targeted applications.

Besides developing applications, there are multiple ways to improve our proposed implementation to make it more scalable and efficient. A trivial enhancement would be the use of static, memory-efficient associative structures, which could present an interesting trade-off for construction index efficiency. We propose to exploit two main characteristics of whole genome sequencing datasets: the existence of an underlying sequence and, therefore, overlapping reads, and the relatively uniform redundancy and error rates. To significantly improve performance beyond exploiting super-*k*-mers, a static index could infer a good ordering (similar to the genome-induced ordering) based on some form of assembly. Using this ordering, we could exploit the fact that many successive *k*-mers in this pseudo-genome would be associated with the same data, improving both query time and index size by reducing redundancy. Furthermore, when not identical, successive *k*-mers often share very similar patterns (see Figure 6b). Therefore, colors themselves could also be encoded relative to the previous colors, as we expect most modifications between two successive colors in the genome to involve the addition or removal of a read. Various degrees of such delta encoding could offer efficient time/memory trade-offs.

**Fig. 6.**
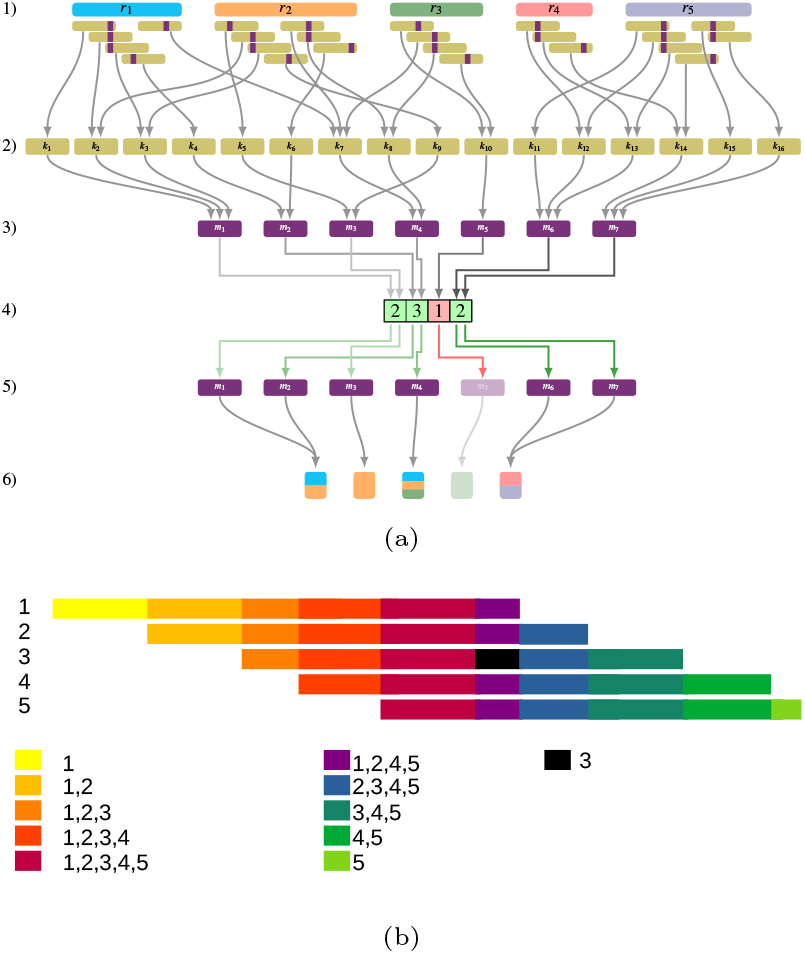
(a) is an example of our lossy version of the Tinted de Bruijn Graph. For a set of reads (1), we can extract the set of *k*-mers (2). Each *k*-mer (2) is linked to a minimizer (3) with possible collisions. By using counting filter (4), we can remove weak minimizer (5). Each remaining minimizer (5) is linked to a color (6). (b) is a simple example of how overlapping reads create subsequences whose *k*-mers are the same colors. We also display the impact of a sequencing errors in the black region that creates read specific sequences.

Optimizing the ordering of reads in sequencing datasets offers a promising approach to significantly enhance the compressibility of list encodings. By reordering the reads in the file to group similar/overlapping reads together, their shared *k*-mers would be associated with highly compressed lists. Such read ordering could be based on clustering, overlap detection steps, or alignment to the aforementioned pseudo-genome. As a proof of concept, we compared the index size constructed from simulated reads in random order and ordered by their origin position in the genome. For a 200X coverage of *C*.*Elegans* with a 1% error rate, the color index size reduced from 3.7GB to 1.3GB, a threefold improvement. With a 0.1% error rate, the color index size goes from 3.4GB to 251MB, an order of magnitude smaller. This display the interest of optimizing read ordering to facilitate their indexability or compression, a problem that start to regain interest for long reads datasets (Lee and Song 2022).

Even though the Minimizer Tinted de Bruijn graph is extremely lightweight and could fit many uses, an actual (*k*-mer) Tinted de Bruijn graph could be interesting for applications where actual de Bruijn graph navigation is required or when a very low number of *k*-mers are queried, and every false positive could be critical. Such an index could mimic the K2R representation and rely on efficient *k*-mer indexes like SSHash (Pibiri 2022).

Additionally, in scenarios where there is a need to search for sequences across multiple datasets, the ability to merge two indexes would be highly beneficial.

Beyond these direct considerations, we believe that the Tinted de Bruijn graph structure is an intriguing representation of a sequencing dataset. Regular de Bruijn graphs are a robust representation of short-read datasets because they encapsulate most of their information in a lightweight and scalable manner. In this manuscript, we demonstrated that a lossy practical implementation is quite lightweight and scalable while retaining almost all the useful information from the original dataset. As a result, while the de Bruijn graph has been deemed unsuitable for noisy long reads, we argue that the Tinted de Bruijn graph is the right representation for such datasets.

We also introduced the concept of the Reversible Tinted de Bruijn graph as a novel form of compressed full-text index adapted to sequencing data. Since we only need to manage duplicated *k*-mers within a given read, we believe that the additional cost to allow a Tinted de Bruijn graph to be reversible will be quite low in most scenarios. The theoretical and practical study of such a structure will be the natural continuation of this work.

## Methods

### Notation and problem

In the following, all the strings will be defined in the DNA’s alphabet (an alphabet of size 4), but all the results could be generalized to a finite alphabet. A *k*-mer of a string *r*, is a substring of *r* of length *k*. For a positive integer *k*, the *k-spectrum* of *r* is the set of all the *k*-mers of *r*. For an integer *m* ≤ *k* and a specific total order of the set of the *m*-mers, the *m-minimizer* (or just minimizer (Roberts et al. 2004)) of a *k*-mer *x*, denoted by *µ*(*x*) is the smallest string of the *k-spectrum* of *x* according to a given order.

To define the set of similar reads *R*^*′*^ ⊆ *R* (where *R* is the initial set of reads) to a sequence *s*, we choose to use the shared *k*-mers number as a filter mimicking (Solomon and Kingsford 2018). A read *r* will then be considered similar to *s* if the number of shared *k*-mers with *s* is greater than a threshold *t*, i.e. |*k*-spectrum(*r*) ∩ *k*-spectrum(*s*)| ≥ *t*.

### Tinted de Bruijn graph

To support multiple queries on the same set of reads, we need to store, for each *k*-mer *x*, the set of reads where it appears, i.e. {*r* ∈ *R* : *x* ∈ *k*-spectrum(*r*)}. As a de Bruijn Graph can be viewed through a set of *k*-mers (by storing arcs, or by testing 4 extensions (alphabet size) to access to possible successive *k*-mers), these two notions are commonly conflated.

Given a document collection *ℱ* (sequencing datasets or assembled genomes), which is a cover for the sets of all reads *R* (∪_*F*∈*ℱ*_ *F* = *R*), a *colored de Bruijn graph* is defined as a de Bruijn graph where each *k*-mer is associated to the documents that contain it, i.e. a mapping from ∪_*r*∈*R*_*k*-spectrum(*r*) to *𝒫*(*ℱ*). *ℱ* is the set of the fasta files, and each *k*-mer is associated with the list of fasta files where it appears.

Similar to the colored de Bruijn Graph, we introduce the *Tinted de Bruijn graph* which is a de Bruijn graph where each *k*-mer is associated to the reads that contain it, i.e. a mapping from ∪_*r*∈*R*_*k*-spectrum(*r*) to *𝒫*(*R*). The Tinted de Bruijn graph can be seen as a special case of colored de Bruijn graph where each read from a sequencing dataset is seen as a distinct source.

We first remark that such structure could, to some extend, recover its original dataset by navigating the de Bruijn graph and reconstructing the reads. However in the case of duplicated region within a given read we can obtain several possible distinct “assemblies” of such reads (multiple Eulerian paths in the induced subgraph of the de Bruijn graph by these reads). This mean that the Tinted de Bruijn graph is not reversible (impossibility to turn up the initial reads and thus some information is lost). Since the presense of repeated *k*-mers corresponds to these non-unique assemblies, by storing, in extra space, the different positions in each read of these *k*-mers, we obtain a reversible version of the *Tinted de Bruijn graph*, which we call the *Reversible Tinted de Bruijn graph*. This reversibility property makes this Reversible Tinted de Bruijn Graph an interesting new element of compressed full-text data structure family, as BWT and FM-index.

Instead of storing the Tinted de Bruijn graph which can be costly, we focus in this work on a lossy version of the Tinted de Bruijn graph, called *Minimizer Tinted de Bruijn graph*, where minimizers are used to represent the *k*-mers as each minimizer *y* is associated with the reads that contains a *k*-mer that has *y* as minimizer, i.e. {*r* ∈ *R* : *y* ∈ {*µ*(*x*) : *x* ∈ *k*-spectrum(*r*)}}.

As for all *k*-mer *x*, {*r* ∈ *R* : *x* ∈ *k*-spectrum(*r*)} ⊆ {*r* ∈ *R* : *µ*(*x*) ∈ {*µ*(*z*) : *z* ∈ *k*-spectrum(*r*)}}, the Minimizer Tinted de Bruijn graph can have false positives but no false negatives. In addition, the scattered positions of the minimizers and the minimizers shared between different *k*-mers mean that this structure is not reversible.

In the following, we present our efficient implementation of the Minimizer Tinted de Bruijn graph dubbed K2R (see Fig. 6a).

### Overview

Given the conceptual similarity between the Tinted de Bruijn graph and the Colored de Bruijn graph, a naive implementation of a Tinted de Bruijn graph would involve an associative structure, such as a hashmap, linking *k*-mers to the list of read identifiers that contain them. Interestingly, an initial tool, SRC (Marchet et al. 2020), developed for short reads, actually implements this strategy using an efficient Minimal Perfect Hash Function (MPHF) (Limasset et al. 2017) as the associative structure. However, we argue that the cost of such a structure would be prohibitively high for large datasets. For example, a human long-reads dataset of 10 million 10kb reads (approximately 33X coverage) would represent more than 3 billion *k*-mers requiring 24GB of storage (for *k* = 31), and each *k*-mer would be associated with dozens of read IDs, necessitating hundred bytes per association. This would mean several hundreds gigabytes of storage without any overhead and without accounting for sequencing errors. In this section, we describe the assumptions we can make when indexing a sequencing dataset and how we can exploit such properties to optimize this naive solution.

### Sparse signal

The primary challenge in scaling colored de Bruijn graphs arises from the inherent difficulty of making a priori assumptions about the *k*-mer distribution within a dataset collection (Almodaresi et al. 2019). In contrast, sequencing datasets predominantly target sequences significantly larger than a kilobase, with relatively low coverage that rarely exceeds a hundredfold. This coverage expectation implies that most *k*-mers will be present in a number of reads closely matching the coverage level, marking their presence in an exceptionally small fraction (by several orders of magnitude) of the total reads, as even bacterial genome sequencing can yield millions of reads. Consequently, unlike with colored de Bruijn graphs, it becomes viable to employ sparse encoding rather than bitvector representations to efficiently denote the presence of *k*-mers in reads.

Employing a 32-bit identifier list to represent *k*-mers turns out to be prohibitively expensive in terms of memory usage, demanding hundreds of bytes per *k*-mer for a standard whole-genome sequencing dataset. A more efficient alternative involves using a sorted list with delta encoding, which significantly reduces the memory footprint. Initially, one might assume that applying compression techniques at such a scale could detrimentally affect construction or query times. However, advancements in compression technology have led to the development of highly optimized algorithms (Trotman and Lin 2016). These algorithms leverage SIMD (Single Instruction, Multiple Data) instructions, enabling near-state-of-the-art compression efficiency with minimal impact on overall throughput. Hereafter, we will refer to a (compressed) list of read identifiers associated with a *k*-mer as a color. As example for a HiFi *E*.*Coli* sequencing with 200X coverage the integers list goes from 1,083MB to 133MB, a 8-fold improvements, without perceptible wall-clock drawback.

### Redundant colors

An important observation in sequencing datasets is the considerable proportion of overlap among reads, which is directly related to the level of coverage. For instance, with 10X coverage, one can anticipate that reads will overlap by approximately 90% of their length, while at 100X coverage, the overlap can increase to 99%, and so forth. In this context, colors in our dataset predominantly represent combinations of reads that overlap, covering specific genomic regions, as illustrated in Figure 6b. Consequently, the diversity of distinct colors observed is relatively limited, since each new read starting position (and consequently, each end position) generates a novel combination. Assuming an ideal scenario devoid of sequencing errors, the number of distinct colors would be expected to increase linearly with the number of reads, necessitating approximately *𝒪*(log(*N*)) bits to encode colors for *N* reads. However, the scenario becomes more complex with the presence of *k*-mers repeated across the genome or *k*-mers that contain sequencing errors. Sequencing errors introduce a significant number of additional colors in two ways: firstly, by creating singleton colors unique to individual reads, and secondly, by generating ‘incomplete’ colors that resemble existing colors but with ‘gaps’ due to the absence of one or more reads, attributable to missing *k*-mers caused by sequencing errors. This leads to the observation that a substantial number of *k*-mers actually share the same color, a property determined by the coverage, read length, and sequencing error rate.

To leverage this characteristic efficiently, we have developed a strategy that indexes only distinct colors by associating them with unique color identifiers, thereby avoiding the redundancy of storing numerous identical colors. This approach entails maintaining a table that links each *k*-mer to its corresponding color identifier, and another table that maps each identifier to the actual list of read identifiers. This method enables the storage of each distinct list precisely once, albeit at the cost of a minor overhead due to the inclusion of identifier integers.

### Successive *k*-mers

Upon closer inspection, we can observe more specifically that overlapping *k*-mers from sequence reads are frequently assigned the same color, primarily because they originate from identical genomic regions and are thus present in the same reads (see Figure 6b). Theoretically, one could attempt to reconstruct such regions and map *k*-mers to their respective regions, which would reflect their shared colors; however, this process is anticipated to be nearly as resource-intensive as genome assembly.

An alternative, more dynamic approach to leverage this characteristic involves the utilization of minimizers. Minimizers, which are the minimal *m*-mers within *k*-mers, are often shared among overlapping *k*-mers (Pibiri et al. 2023), leading to a significantly reduced number of unique minimizers compared to the total number of *k*-mers in a dataset. Ideally, a scheme would select a unique minimizer for every group of *k*−*m k*-mers. Through the application of “random minimizers”, which employ a hashing function, it is estimated that the number of selected minimizers required is twice the minimal theoretical number (Koerkamp and Pibiri 2024). Nevertheless, by adopting advanced minimizer selection algorithms (Zheng et al. 2020; Pellow et al. 2023; Koerkamp and Pibiri 2024), one can surpass these expectations and further reduce the number of selected minimizers in practice.

However, a notable limitation of this technique is the inherent association of all *k*-mers sharing a minimizer with the same color, potentially leading to false positives. To avoid the incidence of false negatives, each minimizer is associated with every read in which it appears. This methodology may result in distinct *k*-mers, which occur in separate reads without shared *k*-mers, logically possessing distinct colors, being assigned the same minimizer and consequently, the same color. Such scenarios, albeit infrequent, could inadvertently introduce false positives, adversely affecting downstream analyses. Conversely, this propensity for false positives may inadvertently counteract the generation of “subcolors” due to sequencing errors, as previously discussed, by incorrectly assigning some erroneous *k*-mers to the correct color, where the genuine genomic *k*-mer would be assigned. We posit that these false positives can be efficiently managed by calculating and reporting the actual number of shared *k*-mers when the selected reads are analyzed, thus providing a balanced approach to minimizing data misinterpretation while maximizing scalability.

### Handling sequencing errors

A fundamental obstacle in addressing our problem is the occurrence of sequencing errors, which introduce novel sequences (and *k*-mers). These are predominantly unique, rendering them incompressible and biologically irrelevant. Moreover, these errors dilute the overall redundancy by transforming abundant sequences into rare novel variants. Traditional full-text indexing approaches often struggle in these scenarios, as they are designed to index entire texts indiscriminately and are typically employed on assembled sequences “cleaned” of sequencing errors for this reason.

In contrast, the success of the de Bruijn graph approach can be partly attributed to its ability to manage sequencing errors effectively by leveraging the concept of *k*-mers abundance. In practice, *k*-mers containing sequencing errors are expected to occur infrequently, whereas the abundance of genomic *k*-mers is closely aligned with sequencing coverage. Thus, implementing a simple abundance filter that retains “solid” *k*-mers appearing more than *T* times in the dataset and excluding “weak” *k*-mers that appear fewer than *T* times can eliminate nearly all erroneous *k*-mers with minimal loss of relevant *k*-mers, provided the threshold *T* is optimally set according to the observed abundance (Chikhi and Medvedev 2014).

Since we focus on indexing minimizers, we introduce a novel variation of this filtering technique applied at the minimizer level. It follows logically that if a *k*-mer is solid (seen more than *T* times), then its minimizer is also considered solid. Consequently, excluding weak minimizers does not result in the exclusion of solid *k*-mers. However, minimizer-level filtering may not effectively eliminate erroneous *k*-mers that share a minimizer with solid *k*-mers. This leads us to anticipate that minimizer filtering is less precise than *k*-mer filtering, with the benefit of significantly reducing the inherent counting cost, as it involves quantifying an order of magnitude fewer keys.

## Data access

We used various datasets : first, several datasets of a small genome (*E*.*Coli*). A recent ONT sequencing with a low error rate (3%), (Accession SRR26899125), a 20kb HiFi sequencing (Accession SRR11434954), and a HiSeq X Ten paired-end sequencing (Accession DRR395239). As a medium-sized genome, we also selected two C.Elegans datasets, one ONT sequencing (Accession SRR24201716) and another HiSeq X Ten paired-end sequencing (Accession ERR10914908). Finally, we selected two human datasets from the T2T project. The HiFi datasets (SRX7897685, SRX7897686, SRX7897687, SRX7897688, and SRX5633451) together amount to 56.8X coverage, comprising both 20kb and 10kb libraries. The ONT sequencings, detailed at http://github.com/marbl/CHM13/blob/master/Sequencing_data.md (126X coverage, error rate of 6%).

## Acknowledgments

This work is funded the French National Research Agency AGATE ANR-21-CE45-0012.

