## Appendix for "K2R: Tinted de Bruijn Graphs implementation for efficient read extraction from sequencing datasets"

### 1 Implementation Details

#### 1.1 Index Structure

In our index structure, we employ a dual mapping system consisting of a [minimizer: color identifier] map and an [color identifier: color] map, where minimizers and color identifiers are integers, and colors are an object composed of a compressed integer array, an integer array and several integers (representing structure sizes and occurrencies, for implementation purpose). W e opted to use the unordered_dense hash map, available at `http://github.com/martinus/unordered_dense`, due to its exceptional performance characteristics, which have been leveraged in several bioinformatics tools, including kallisto (Bray et al., 2016) and mashmap (Jain et al., 2017).

Upon inserting a read into the index, its minimizers are computed and inserted into the minimizer map. Each corresponding identifier is then updated to incorporate the new read identifier into its color list. If the associated color does not preexist within the map, a new identifier is generated and added to the identifier map, along with its newly associated color. Colors are compressed using the TurboPFor library `http://github.com/powturbo/TurboPFor-Integer-Compression`, which displays almost state-of-the-art compression levels at a very low computational cost.

To keep the index lean and efficient, the identifier map also tracks the number of minimizers referencing each identifier. When a minimizer's association shifts to a different identifier, two scenarios arise. First, if one or more minimizers is still associated to the former color identifier, if the new color exists, the minimizer can be automatically associated to it, otherwise a new color is created using the former one. Second, if the color is no more associated to any minimizer, this color is not deleted but modified to correspond to its last minimizers, adding a read identifier.

Our choice of index structure imbues it with dynamic capabilities, allowing for real-time querying at any point during the read insertion process. While static structures may offer advantages in memory efficiency, they lack flexibility and require a more cumbersome construction phase.

#### 1.2 Minimizer Abundance Filtering

To enhance construction efficiency and minimize the index size, we implement an optional filtering mechanism for weak minimizers using a Counting Bloom filter to approximate minimizer abundance within the dataset. It's important to note that a Counting Bloom filter can only overestimate counts, leading to false positives by misclassifying some weak minimizers as solid due to hash collisions with genuinely solid minimizers.

After approximate minimizer counting, we convert the Counting Bloom filter into a Regular Bloom filter by transforming each cell, originally representing counts with integers, into a single bit. This bit indicates the solidity of the associated minimizer, significantly reducing memory consumption during the actual index construction. Using this strategy, we limit the peak memory usage as the hash tables should become the memory bottleneck using a well sized Bloom filter.

We also observe the opposite pattern of overabundant minimizers linked to highly repeated regions in a genome, either due to biological mechanisms or sequencing type. Since such minimizers can be associated with an incredibly high number of reads, they can seriously hinder index performance and "pollute" the output with many irrelevant matches. To counteract this, we implement an optional maximum minimizer abundance filtering, as commonly performed by other tools (Li, 2018).

When employing minimizers in place of $k$-mers, alongside Counting Bloom filters to sift through them, a natural concern arises regarding the potential influence of false positives introduced by this method on the final output. To address this, we conduct an evaluation illustrated in Figure S1 of the Appendix, where we calculate the proportion of reads selected based on their minimizer content relative to the total number of reads achieving the requisite $k$-mer similarity threshold (here fixed at 0.3). Remarkably, a substantial majority of queries exhibit a ratio exceeding 75%, indicating that a majority of the selected reads are, in fact, akin to the query. This high ratio underscores the efficiency of our approach in maintaining a strong correlation between selected reads and their relevance to the query.

#### 1.3 Parallelization Overview

The filtering phase is parallelized with a mutex array to secure sections of the Counting Bloom Filter, enabling efficient concurrent operations. Parallelizing the index construction phase is more complex due to dependencies on synchronized maps. We mitigate this by implementing an inter-reads parallelization, where minimizers are computed across sequence substrings concurrently. Tasks related to color management, including decompression, updates, sort and recompression, are executed in parallel, protected by mutex arrays to ensure data integrity. This setup allows concurrent modifications to the color and $m$-mer maps without data corruption.

The queries can also be executed in parallel, with each thread handling a single query.

#### 1.4 Minimizer Scheme

Minimizers reduce complexity by representing a set of overlapping $k$-mers with a single $m$-mer. The ideal minimizer selection scheme would identify one $m$-mer for every $k-m+1$ sequence of overlapping $k$-mers. Practical implementations, such as the random minimizer strategy, tend to select approximately twice as many minimizers as the ideal case.

Recent advancements in minimizer selection techniques aim to closely approach this theoretical lower bound, thus reducing the quantity of necessary minimizers. We employ decycling set minimizers (Pellow et al., 2023), which minimize the count of selected minimizers, albeit at the cost of increased computational overhead.

##### 1.5 Homocompression

To address homopolymer run length errors prevalent in HiFi and PacBio sequencing reads, we propose an optional 'homocompression' feature. This technique lossily represents sequences within reads by compressing consecutive occurrences of a nucleotide X...X into a single instance of X. As highlighted in the La Jolla Assembler paper (Bankevich et al., 2022), applying homocompression to HiFi reads can result in a threefold reduction in total error count, making a substantial proportion of reads error-free. This approach not only improves sequencing data accuracy but also enhances the overall reliability of genomic analysis by significantly reducing the impact of homopolymer-associated errors.

#### 2 Supplementary Figures

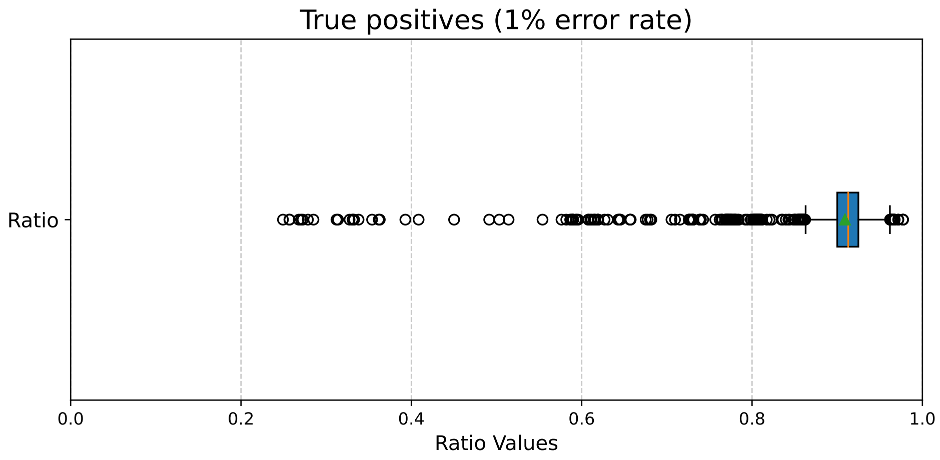

(a)

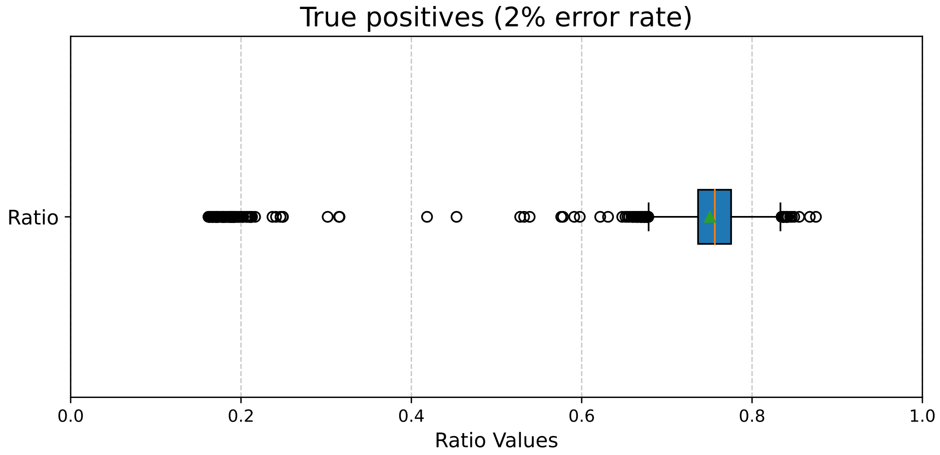

(b)

Figure S1: Ratio between verified and kept reads, during queries of 10,000 sequences of length 10,000 without any errors using K2R with a threshold of 0.3. The index is created from simulated reads from the *E.Coli* genome of length 10,000, with a 200X coverage and an error rate of 1% in (a) and 2% in (b).

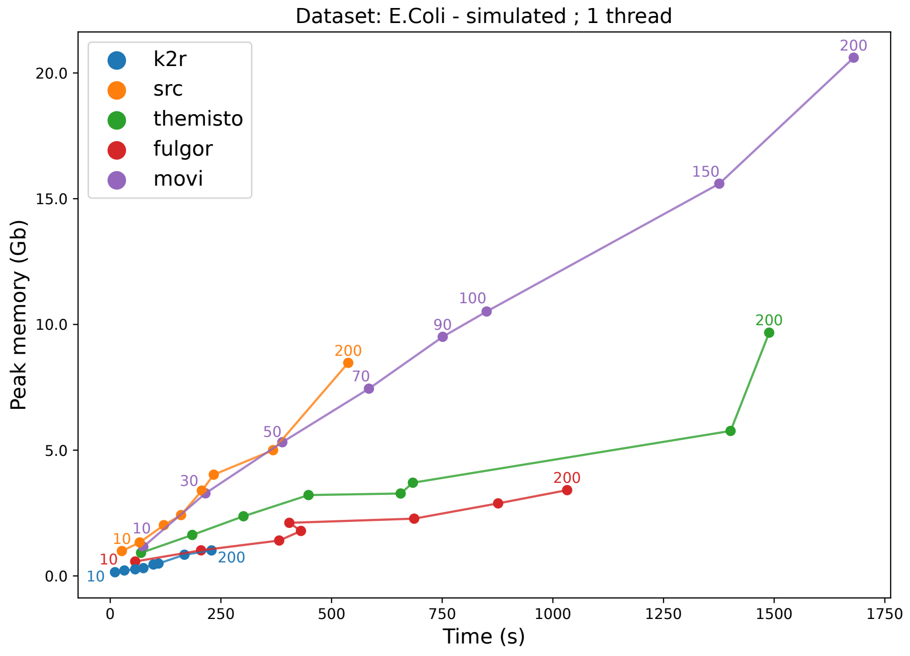

(a)

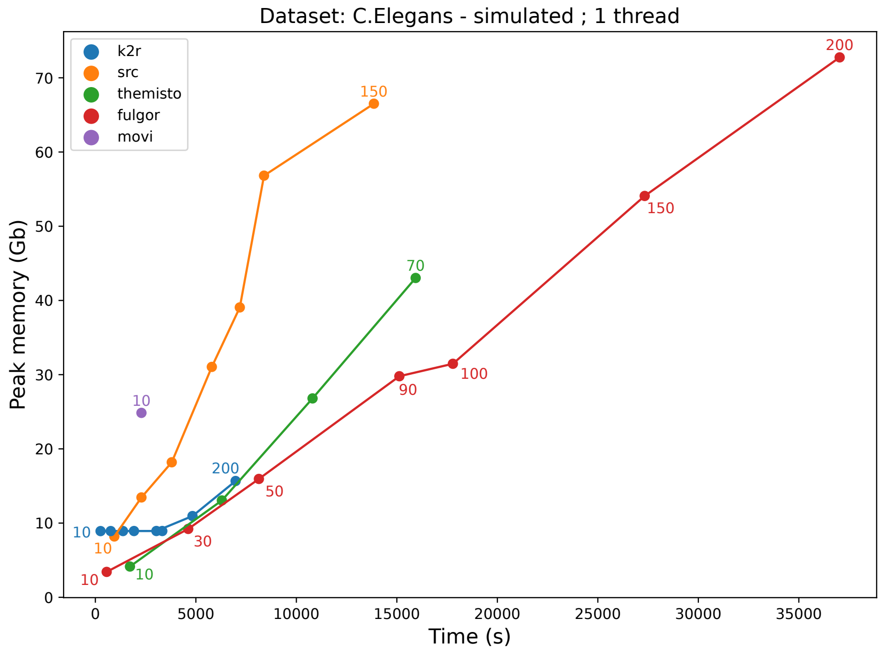

(b)

Figure S2: Comparison of ressource usage (wall-clock time and memory peak) used during the index construction with K2R, SRC, Movi, Fulgor and Themisto according to the available coverages. The dataset used consists of simulated reads from (a) *E.Coli* and (b) *C.Elegans* genomes, each with a length of 10,000 and a 1% error rate.

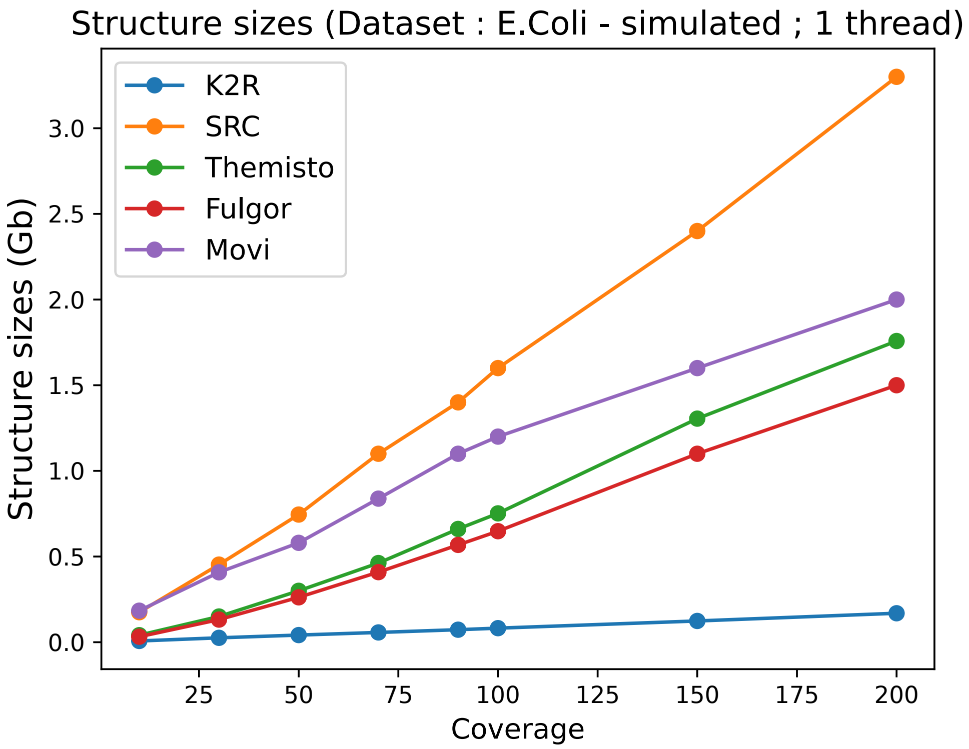

(a)

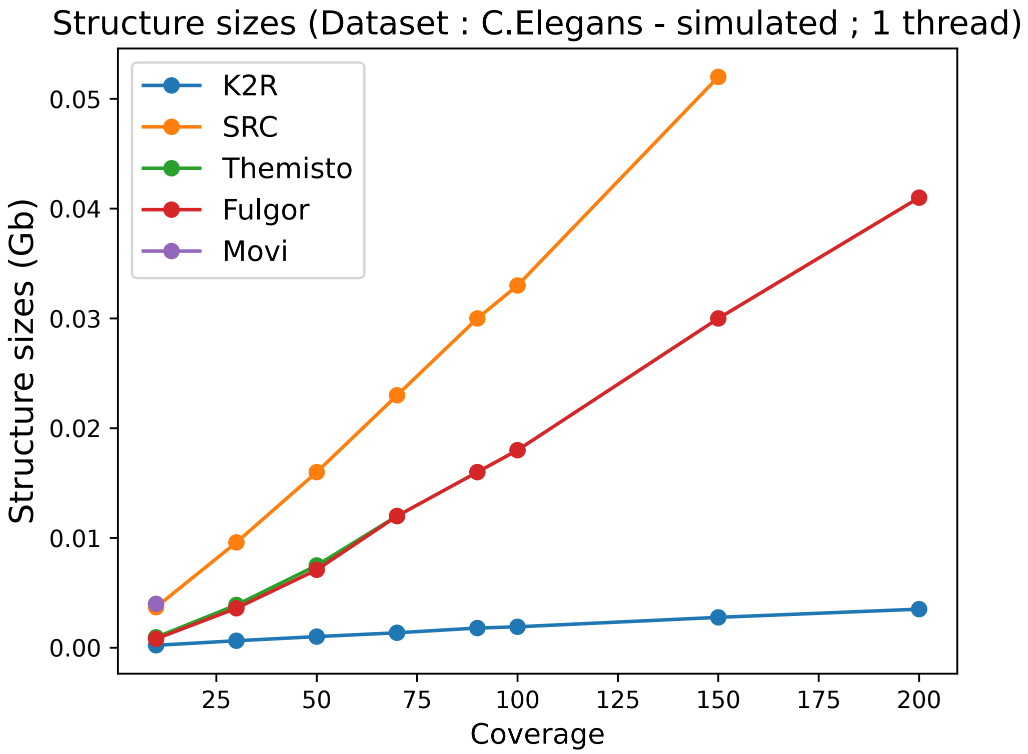

(b)

Figure S3: Comparison of the index size for different coverage values using simulated reads from (a) *E.Coli* and (b) *C.Elegans* reference genomes, with a 1% error rate and a read length of 10,000.

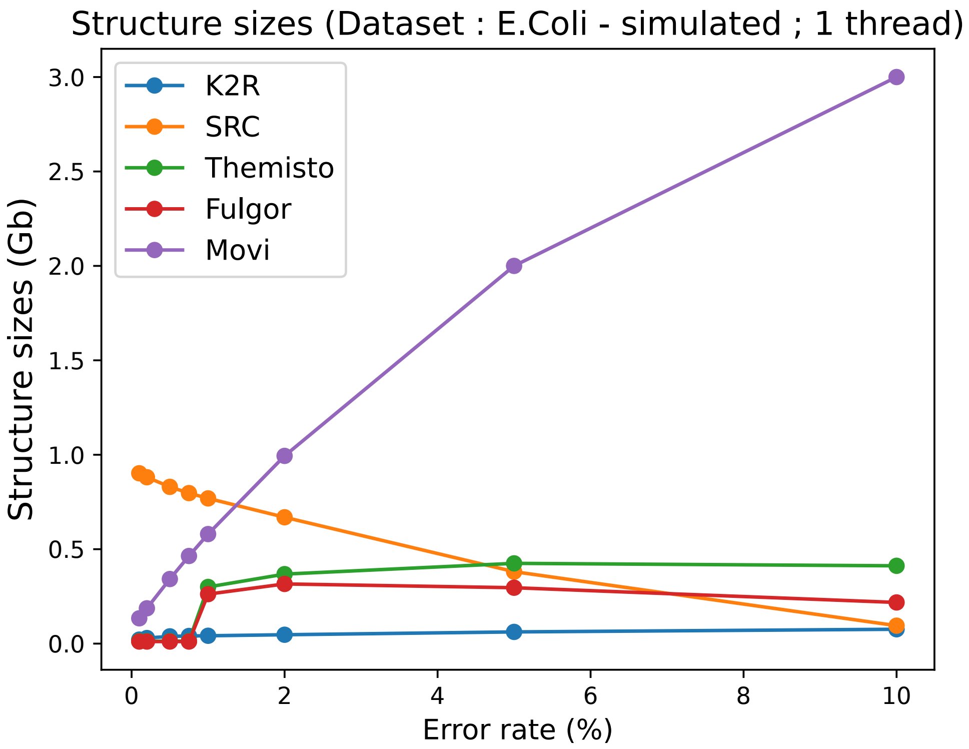

(a)

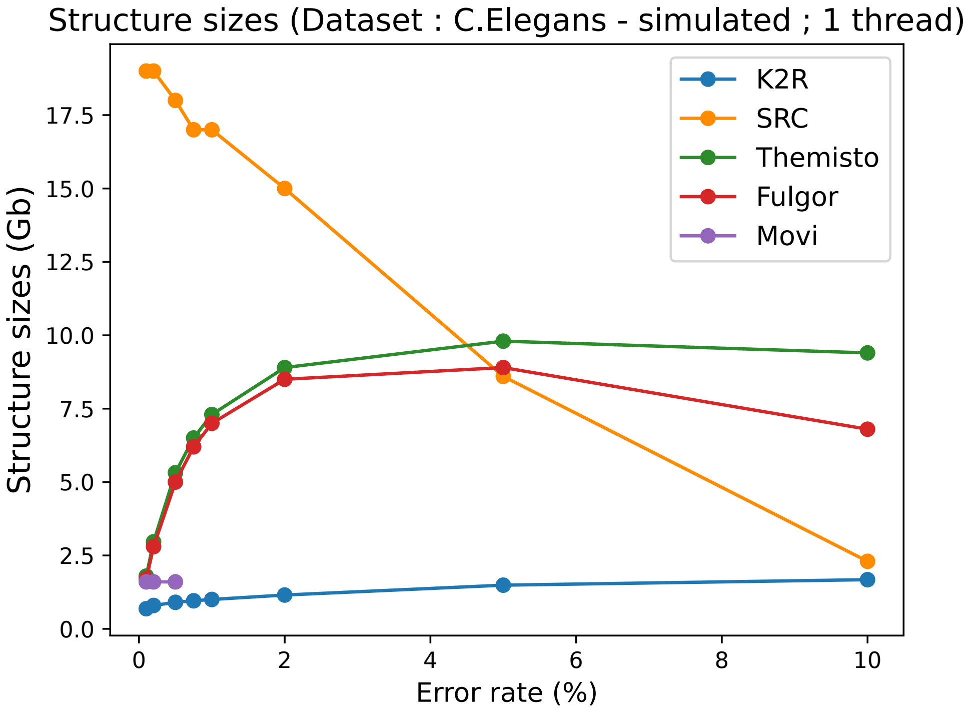

(b)

Figure S4: Comparison of the index size for different error rate values using simulated reads from (a) *E.Coli* and (b) *C.Elegans* reference genomes, with a coverage of 50X and a read length of 10,000.

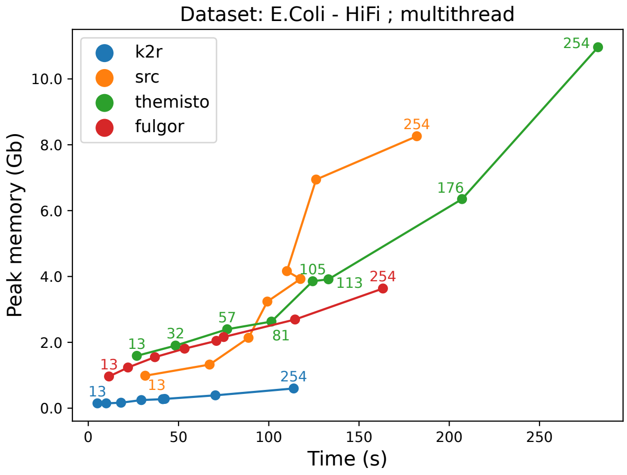

(a)

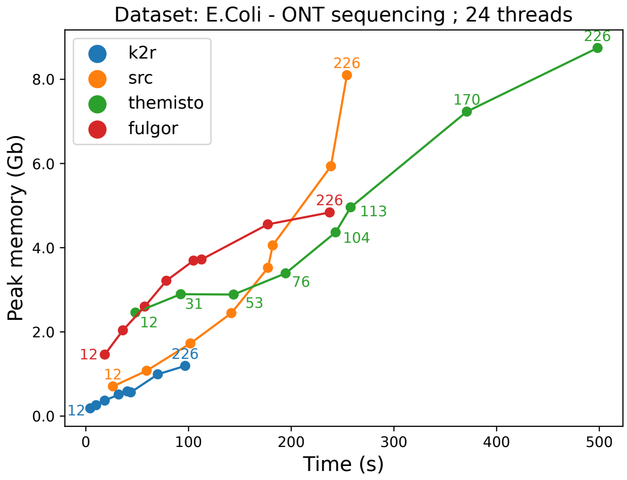

(b)

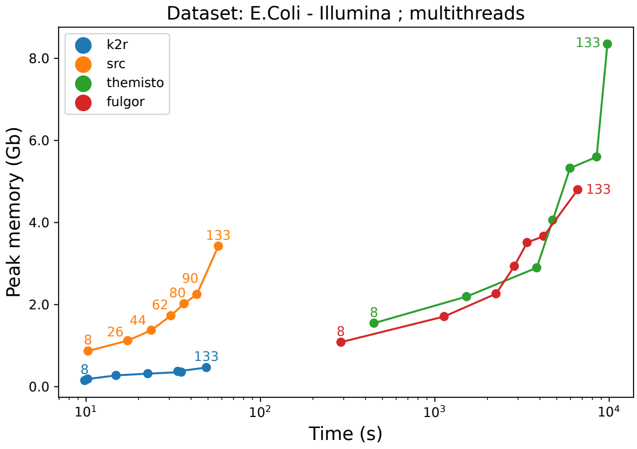

(c)

Figure S5: Comparison of ressource usage (wall-clock time and memory peak) used during the index construction with K2R, SRC, Fulgor and Themisto according to the available coverages, with several threads. The dataset used consists of reads from 3 different datasets of *E.Coli* genome : (a) HiFi dataset, (b) ONT dataset and (c) Illumina dataset, with different coverages.

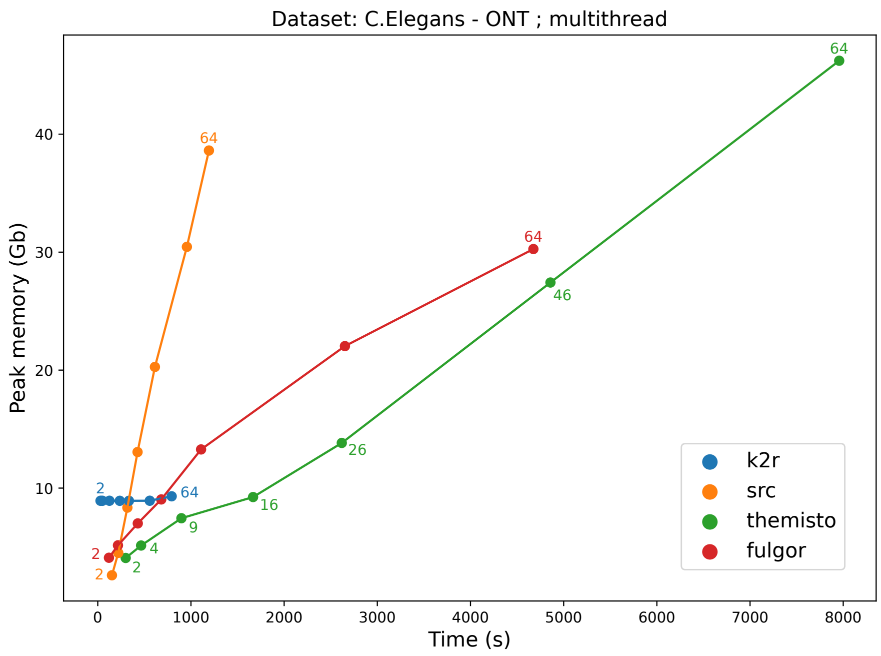

(a)

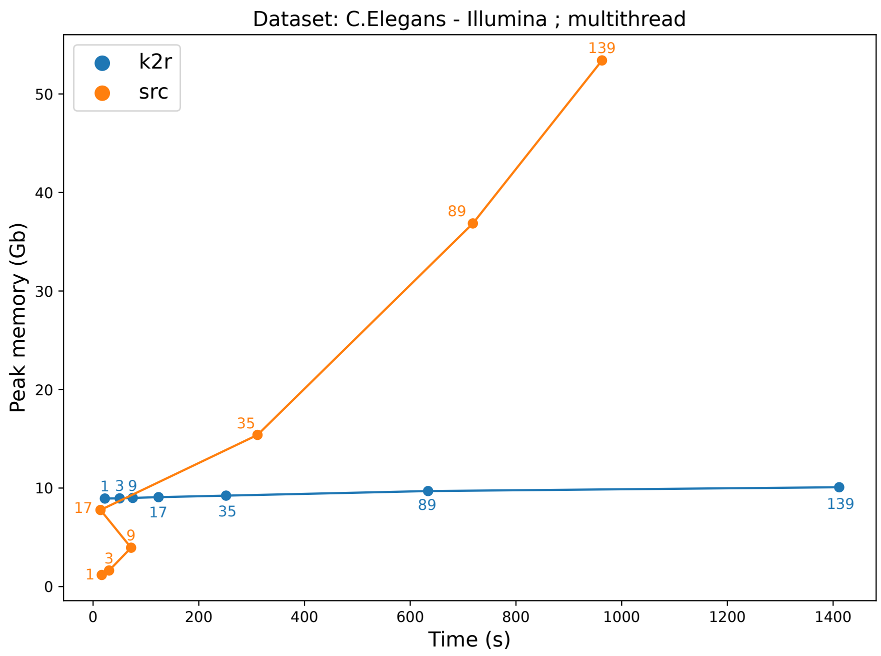

(b)

Figure S6: Comparison of ressource usage (wall-clock time and memory peak) used during the index construction with K2R and SRC, according to the available coverages, with several threads. The datasets used consists of reads from 2 different datasets of *C.Elegans* genome : (a) ONT dataset and (b) Illumina dataset, with different coverages. We notice that Themisto, Fulgor and Movi can't be tested here, because of their mode of use and scale up problems respectively.

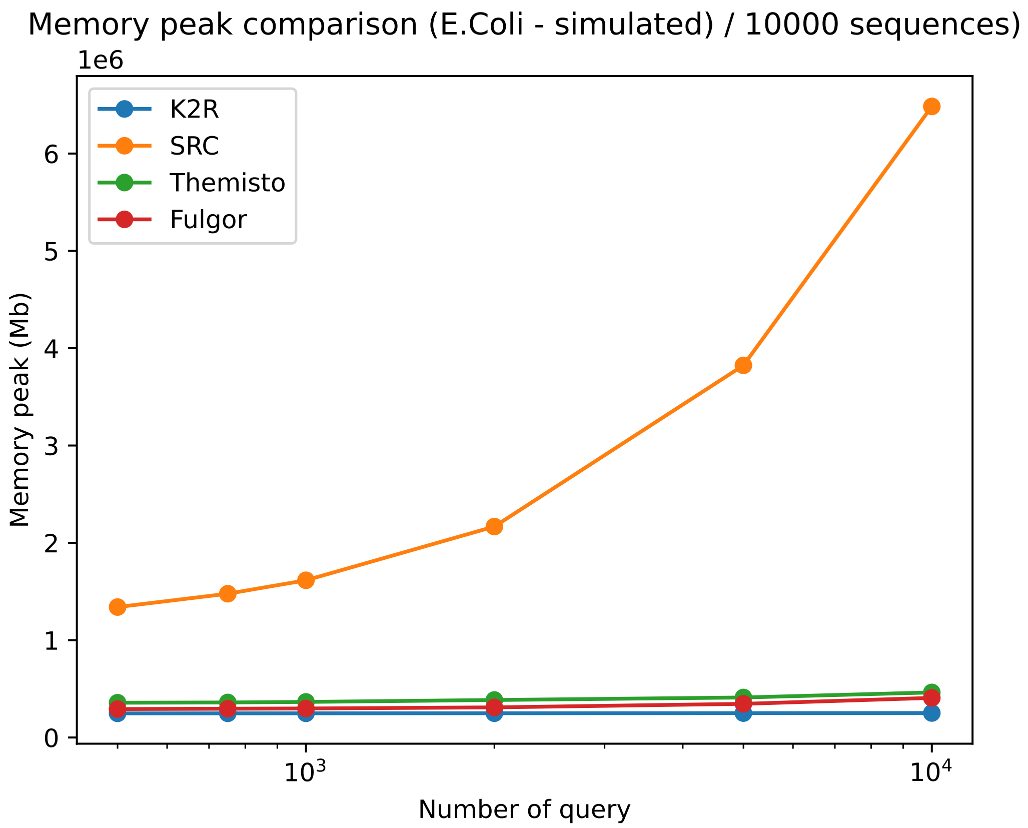

(a)

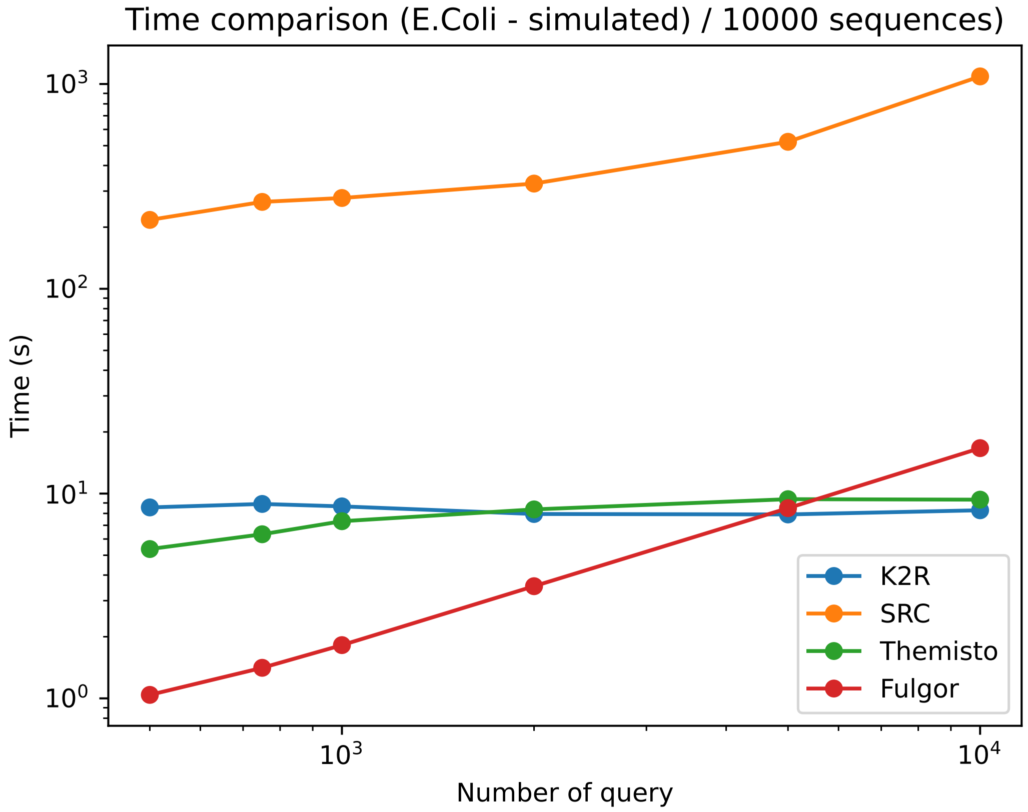

(b)

Figure S7: Comparison of the memory peak (a) and time consuming (b) during ”positive” queries with K2R, SRC, Fulgor and Themisto according to the number of queried sequences. The sequences queried consists of 10,000 reads of length 10,000 from a simulated *E.Coli* dataset, having a 50X coverage and a 1% error rate.

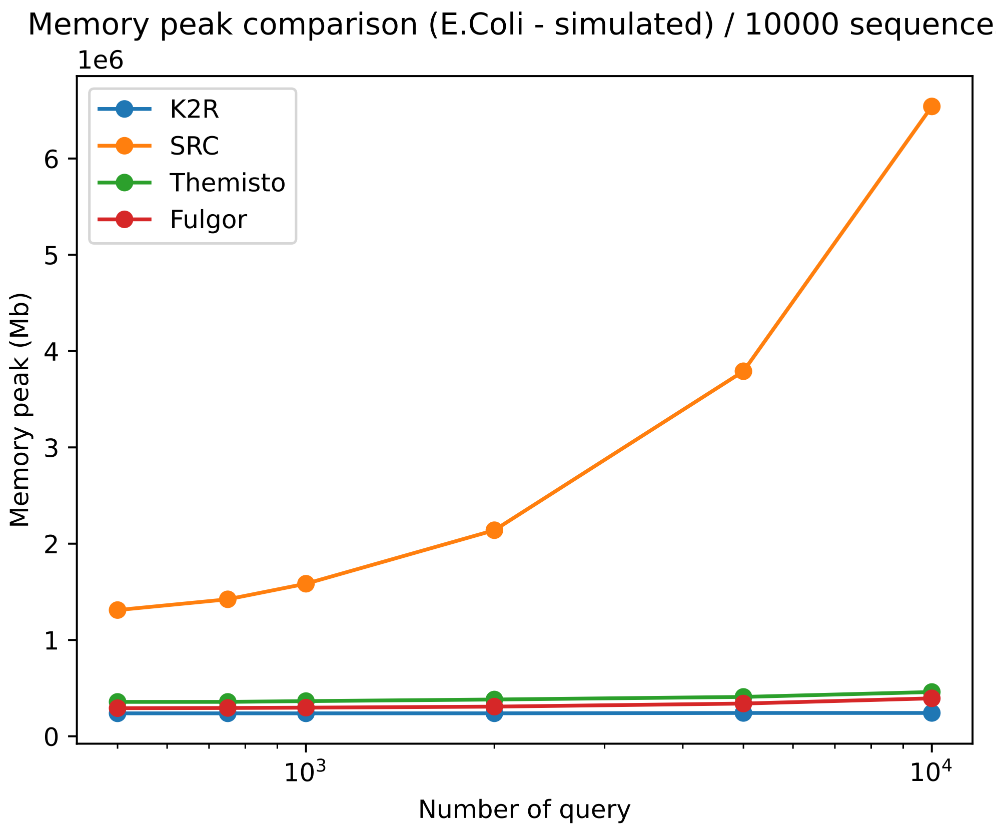

(a)

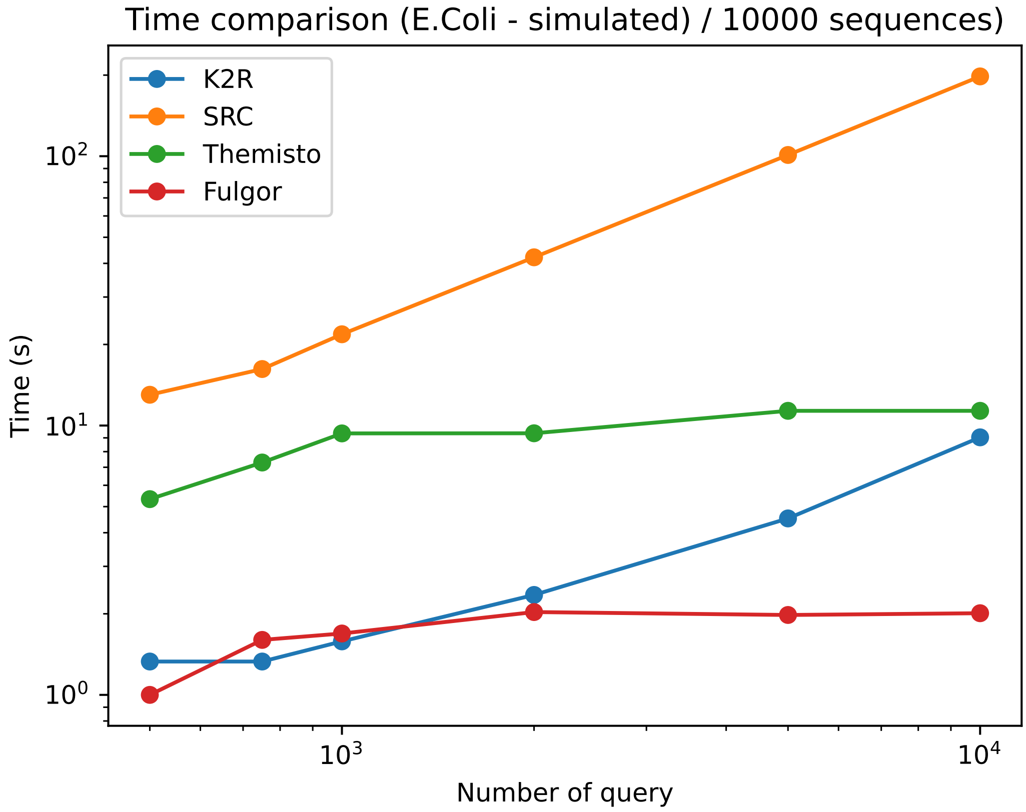

(b)

Figure S8: Comparison of the memory peak (a) and time consuming (b) during ”negative” queries with K2R, SRC, Fulgor and Themisto according to the number of queried sequences. The sequences queried consists of 10,000 reads of length 10,000 from a simulated *E.Coli* dataset, having a 50X coverage and a 1% error rate.

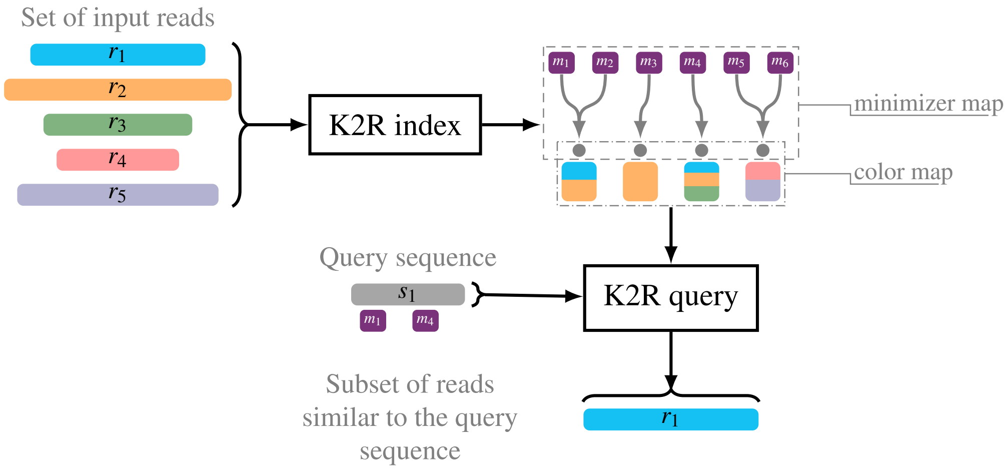

(a) Global use case of K2R: for finding the subset of reads similar to a sequence, we first use `K2R index` to index all the available reads. In a second step, we use `K2R query` on the sequence to find similar reads by using the index.

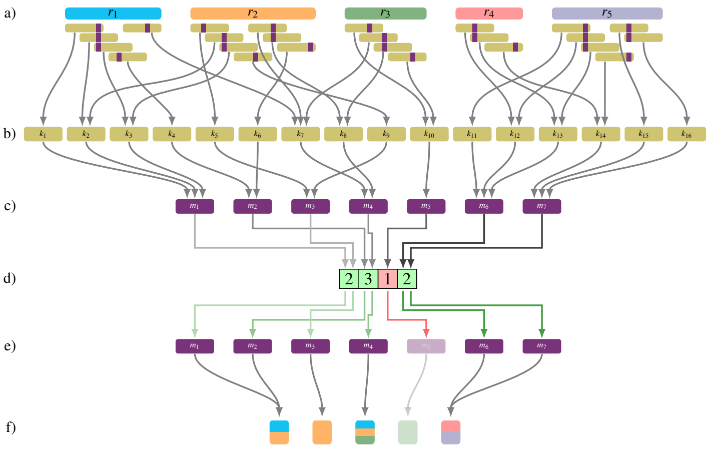

(b) `K2R index`: For a set of input reads (a), we can extract the set of $k$-mers (b). Each $k$-mer (b) is linked to a minimizer (c) with possible collisions. By using counting filter (d), we can remove weak minimizer (e). Each remaining minimizer (e) is linked to a color (f). We store this link from minimizers (e) to colors (f) by two maps : the *minimizer map* and the *color map*.

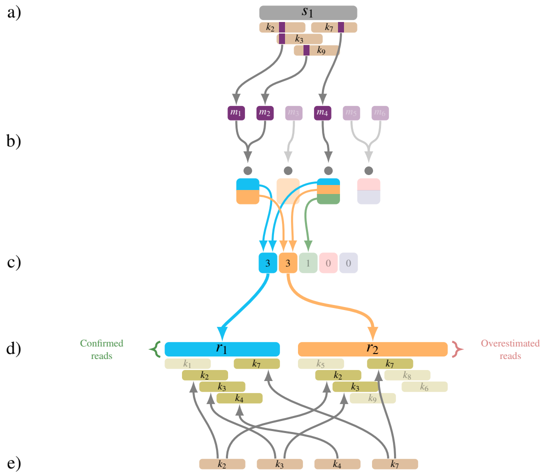

(c) `K2R query`: For a query sequence (a), we extract the set of $k$-mers and minimizers. We use each minmizer of the query sequence in the index constructed by `K2R index` (b) to compute the counting table of the reads indexed (c) to select potential similar reads (d). Each $k$-mer of the query sequence is searched for in each of the potential similar reads (e) to remove overestimated reads.

Figure S9: `K2R`
